# Generation of star-shaped human induced astrocytes with a mature inflammatory phenotype for CNS disease modeling

**DOI:** 10.1101/2021.06.07.447336

**Authors:** Dimitrios Voulgaris, Polyxeni Nikolakolpoulou, Anna Herland

## Abstract

Generating astrocytes from induced pluripotent stem cells has been hampered by either prolonged differentiation -spanning over two months -or by shorter protocols that generate immature astrocytes, devoid of salient inflammation-associated astrocytic traits pivotal for CNS neuropathological modeling. We directed human neural stem cells derived from induced pluripotent stem cells to astrocytic commitment and maturity by orchestrating an astrocytic-tuned culturing environment. In under 28 days, the generated cells express canonical and mature astrocytic markers, denoted by the expression of AQP4 and, remarkably, the expression and functionality of glutamate transporter EAAT2. We also show that this protocol generates astrocytes that encompass traits critical in CNS disease modeling, such as glutathione synthesis and secretion, upregulation of ICAM-1 and a cytokine secretion profile which is on par with primary astrocytes. This protocol generates a multifaceted astrocytic model suitable for CNS in vitro disease modeling and personalized medicine through brain-on-chip technologies.

## Introduction

Astrocytes have long been considered as merely cellular scaffolds - i.e., the brain’s glue. Recent studies point out that astrocytes have a prominent role in health and disease; once considered a recycler of neurotransmitters and a supply cabinet for neurons, this star-shaped cell type has an ever-growing role in health and disease (Araque *et al*., 1999; Barres, 2008; Liddelow *et al*., 2017).

Astrocytes are a truly versatile brain cell type, regulating a plethora of cellular processes such as glutamate clearance, provision of antioxidants to neurons and relay of inflammatory signals. Additionally, astrocytes actively participate and regulate synaptic transmission (Araque *et al*., 1999; Perea, Navarrete and Araque, 2009), and they are involved in the pathophysiology of numerous degenerative diseases (Liddelow *et al*., 2017).

Importantly, mouse and human astrocytes are transcriptionally and functionally different (Zhang *et al*., 2016), human astrocytes are larger and have more elaborate processes than mouse astrocytes (Oberheim *et al*., 2009). Another notable difference is the expression of glutamate transporters EAAT1 and EAAT2; in mouse, GLAST (the mouse equivalent of EAAT1) expression precedes GLT-1 (the mouse equivalent of EAAT2), the latter denotes a postnatal phenotype. Conversely, both transporters are expressed prenatally in humans(DeSilva *et al*., 2012, p. 2). Additionally, pathological conditions, such as multiple sclerosis (MS), affect humans exclusively(‘t Hart, 2016) and experimentally induced mouse models fail to deepen our knowledge in potential treatments (Sriram and Steiner, 2005). Stem cell technology paved the way for the generation of numerous differentiation protocols that generate human induced astrocytes (hiAstrocytes), bypassing the need for primary sources, and bringing forth human models that can potentially capture in higher biofidelity human astrocytes than mouse models.

Astrocytes respond to pathological conditions in a disease-specific fashion; in ALS, the glutamate transporter EAAT2 is lost in astrocytes suggesting a causal relationship between neuronal excitotoxicity and EAAT2 loss in astrocytes(Rothstein *et al*., 1995). Apart from inflammatory cytokine secretion upon stimulation (e.g., IL-6 and IL-8 (Choi *et al*., 2014)), astrocytes can also express ICAM-1 in pathological conditions (Frohman *et al*., 1989) such as MS(Brosnan *et al*., 1995), brain injury and AD (Müller, 2019). In MS, ICAM-1 on astrocytes serves as direct communication with infiltrating leukocytes (Héry *et al*., 1995; Lee *et al*., 2000, p. 1) and microglia(Akiyama *et al*., 1993). Astrocytes are also unique shapeshifters; they remodel their processes and orient them towards lesions (Schiweck, Eickholt and Murk, 2018) while in transgenic AD animals, astrocytes exhibit reduced complexity in their processes (Rodríguez *et al*., 2009). In epilepsy, astrocytic processes appear thicker and longer (Oberheim *et al*., 2008). Astrocytes synthesize and secrete glutathione (GSH)(Yudkoff *et al*., 1990; Sagara, Makino and Bannai, 1996) shielding CNS cells from oxidative stress, which is also reflected in the high astrocytic GSH content ~8mM (Dringen and Hamprecht, 1998). Interestingly, GSH availability is altered in TBI (Harris *et al*., 2012) and autism(Rose *et al*., 2012; Gu *et al*., 2013), while GSH efflux is impaired in AD(Lovell, Xie and Markesbery, 1998; Sultana and Butterfield, 2004). It remains to be determined whether GSH disturbance is the etiology, contributing factor or simply an effect of pathological conditions. Arguably, astrocytes have indeed a prominent role in pathological conditions of the CNS.

The need to model interactions of the CNS necessitates reliable and sustainable (i.e., non-primary) sources of human astrocytes. The extent of information extracted from an *in vitro* study is limited by the model used; hence, a holistic view of astrocytes in pathology demands the concurrent existence of a spectrum of traits that exceed the current traits of astrocytic differentiations. To recapitulate pathological conditions *in vitro*, an astrocytic model, among others, should: 1) express and have functionally active EAAT1 and EAAT2 transporters 2) capacity for an immune response upon inflammatory stimuli both via cytokine secretion and upregulation of inflammatory adhesion markers (i.e., ICAM-1) that is comparable to primary astrocytes and 3) synthesize and secrete GSH that is comparable to primary astrocytes.

There is a lack of a short astrocytic differentiation protocol (<30 days) from a stable precursor that can recapitulate all the mentioned processes. Currently, astrocyte generation is hampered by either extensive differentiations (>60 days) (Roybon *et al*., 2013; Holmqvist *et al*., 2015; Oksanen *et al*., 2017, p. 1; Perriot *et al*., 2018) or by shortened differentiation protocols (<60 days) that lack a process-bearing phenotype and GSH synthesis/secretion (Santos *et al*., 2017). Our recent efforts in astrocyte generation yielded GFAP-negative cells that lack EAAT2 functionality, thus resembling an immature phenotype (Lundin *et al*., 2018, 2020). We envisioned that we could generate mature astrocytes with *in vivo*-like morphology and functionality by providing an astrocytic-tuned milieu to neural stem cells. An astrocytogenic milieu comprises a suitable differentiation media, an astrocytic-tuned ECM coating and appropriate cell-to-cell contact (Li *et al*., 2019).

Almost all astrocyte differentiations in a monolayer format have been carried out in coatings such as Matrigel or laminins. However, none of these ECM molecules are used for *in vitro* culturing of human primary astrocytes. An interesting candidate ECM molecule is collagen; collagen is indeed not abundant in the bulk ECM of the CNS (Zimmermann and Dours-Zimmermann, 2008) and is mainly restricted in the basement membrane and meninges. In vitro studies unveiled that astrocytes express fibrillar collagen, which is inhibited *in vivo* by EGF signaling and meningeal cells (Heck *et al*., 2003, 2007). Interestingly, when fetal astrocytes were cultured in a 3D collagen gel, they assumed a star-like morphology instead of the elongated phenotype that is ubiquitous in a conventional culturing environment (Placone *et al*., 2015). As opposed to laminins, collagen is not well suited for neuronal differentiations (Ma *et al*., 2008), hence creating an ECM environment that can potentially boost astrocytic commitment and dampen the neurogenic potential of neural stem cells. Gelatin, a denatured form of collagen, has been used for the primary isolation of astrocytes(Sinyuk and Williams, 2020); thus we elected to use it in our differentiation strategy.

Here, we report on a differentiation strategy that unleashes the astrocytic potential of iPS-derived neural stem cells without any shorting steps. By day 28, human iPS-derived astrocytes (hiAstrocytes) feature a star-shaped morphology, display inflammatory potency and functional uptake of both astrocytic glutamate transporters (EAAT1 and EAAT2) as well as an mRNA and protein profile that resembles human astrocytes. Additionally, we demonstrate that hiAstrocytes synthesize and secrete glutathione. This astrocytic model can be used for CNS disease modeling since it encompasses a multitude of traits that are altered during pathological conditions.

## Results

### Neural stem cells generate star-shaped cells in 28 days under astrocytogenic conditions

Neuroepithelial cells (NES) are an intermediate cellular stage derived from induced pluripotent stem cells that can generate both neurons and astrocytes. NES can be cryopreserved and cultured up to 100 passages without major phenotype changes and are therefore a robust starting point for neural differentiation(Falk *et al*., 2012). We used three iPS-derived neural stem cell lines, NES C9, NES C7 and NES AF22. Two of these lines (NES C9 and NES AF22) lines have been previously used for astrocyte differentiation (Lundin *et al*., 2018, 2020). Here, we differentiated the neural stem cells for 28 days (Figure 1a) using a primary astrocytic (AM) media that has been previously shown to induce astrocytic traits in various research groups (Tcw *et al*., 2017; Soubannier *et al*., 2020). In this approach, the growth factor cocktail is tailored for astrocytic growth and maintenance (Michler-Stuke, Wolff and Bottenstein, 1984; Haselbacher *et al*., 1989; Codeluppi *et al*., 2011).

**Figure 1.**
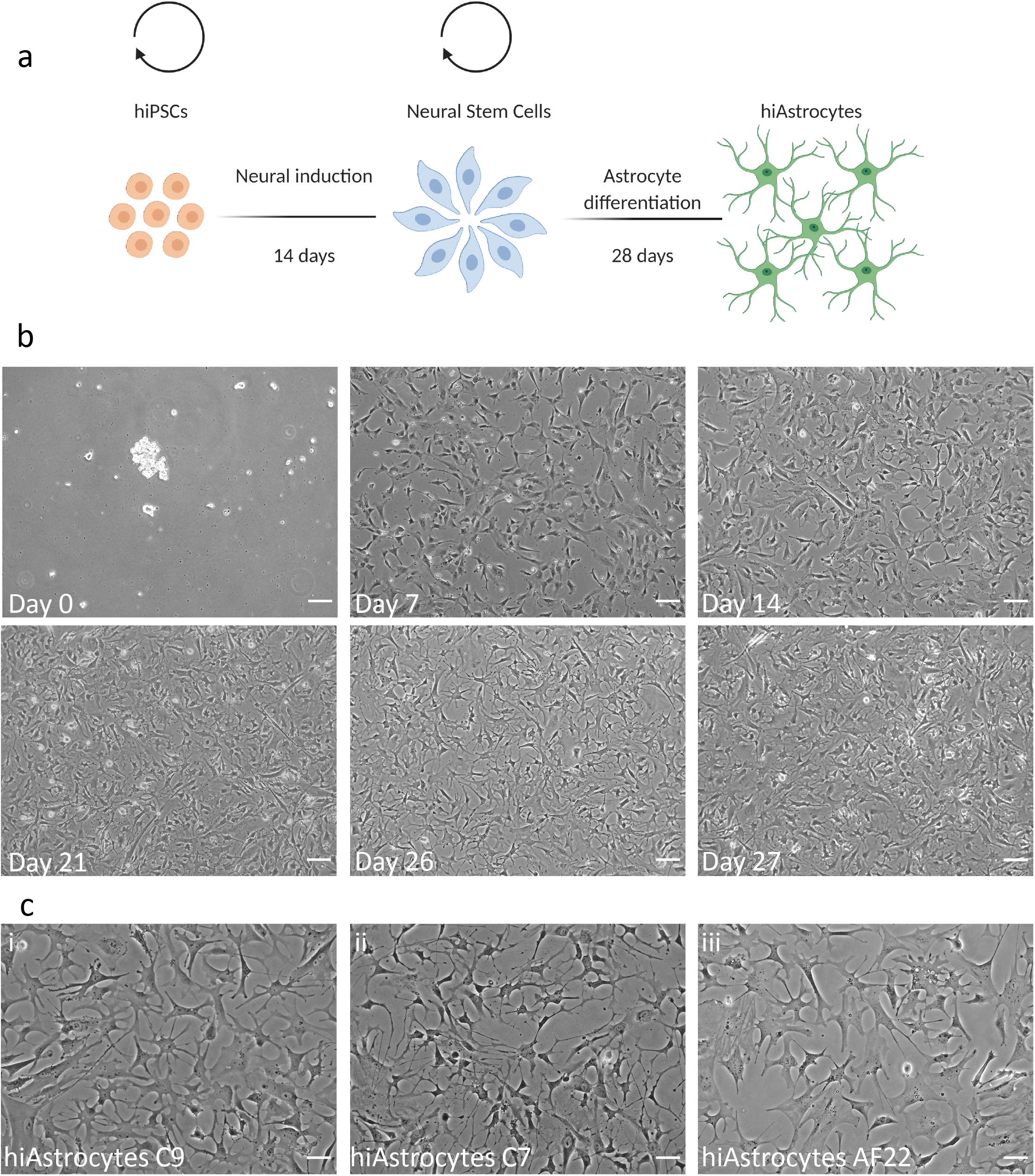
Neural stem cells generate star-shaped cells in 28 days under astrocytogenic conditions. a) Schematic presentation of the duration of the protocols used to derive neuroepithelial stem cells and hiAstrocytes. b) Brightfield images of NES C9 for differentiation days 0,7,14, 21, 26 and 27. Scale bar 100 μm. c) Brightfield images of hiAstrocytes C9. (i) hiAstrocytes C7 (ii) and hiAstrocytes AF22 (iii) on day 26 of differentiation. Scale bar 50 μm.

Neural stem cells were transiently cultured in suspension as aggregates and further differentiated in a monolayer format. Cells were seeded at 288K/well in a 6-well plate format (day −1) in 0.2% (w/v) gelatin-coated wells in N2B27 media. The day after (Figure 1b, day 0, NES C9), most cells were in suspension, forming small aggregates. Media was carefully removed, and cells were washed once with DPBS (w/ Ca^++^ and Mg^++^) before adding AM media supplemented with AGS and FBS.

The day after (Figure 1b, day 0), the cell aggregates had attached to the culture vessels, and we passaged (at 30K/cm2) the cultures after 1-2 days upon reaching confluency since that is pivotal for the morphology of hiAstrocytes. The first 6 days are crucial for a successful astrocyte differentiation; cells assume a colony-like morphology, growing outwards. By day 7, cells lost their NES-like behavior; namely, cells did not grow in NES colonies upon passaging, appeared more elongated, and a small percentage of cells assumed a triangular morphology (Figure 1b, day 7). By day 14, the frequency of triangular-shaped cells increased (Figure 1b, day 14) while the growth rate decreased (compared to NES), suggesting a switch to differentiation over self-replication. By day 21, cellular processes could be detected emanating from the somata, resembling star-shaped morphologies. The proliferation rate was further reduced by day 26 (Figure 1b, day 26), and cells transitioned to more noticeable morphological changes assuming a star-like morphology (Figure 1b, day 27), apparent in all lines (Figure 1c, i) NES C9 ii) NES C7 and iii) AF22).

### HiAstrocytes have a distinct astrocytic mRNA expression profile that differs from spontaneously differentiated cells

We characterized hiAstrocytes by RT-qPCR and compared them to human fetal cortical astrocytes (HFA) at low passage (p.3). In hiAstrocytes C9, the so-called gliogenic switch, Nuclear factor IA (NFIA)(Tchieu *et al*., 2019), was upregulated during the differentiation while the neuronal progenitor marker *DCX* was downregulated (Figure 2a), denoting glial commitment. Most astrocytic markers such as *CD44, GFAP* and *ALDH1L1* were on par with the expression levels in HFA, while *AQP4* and *S100B* were significantly enriched in hiAstrocytes opposed to HFA (Figure 2b). Interestingly, NES and HFA had the same expression level of the astrocytic marker S100B. HiAstrocytes C7 and AF22 had an expression pattern that was comparable to hiAstrocytes C9 (Figure 2c).

**Figure 2.**
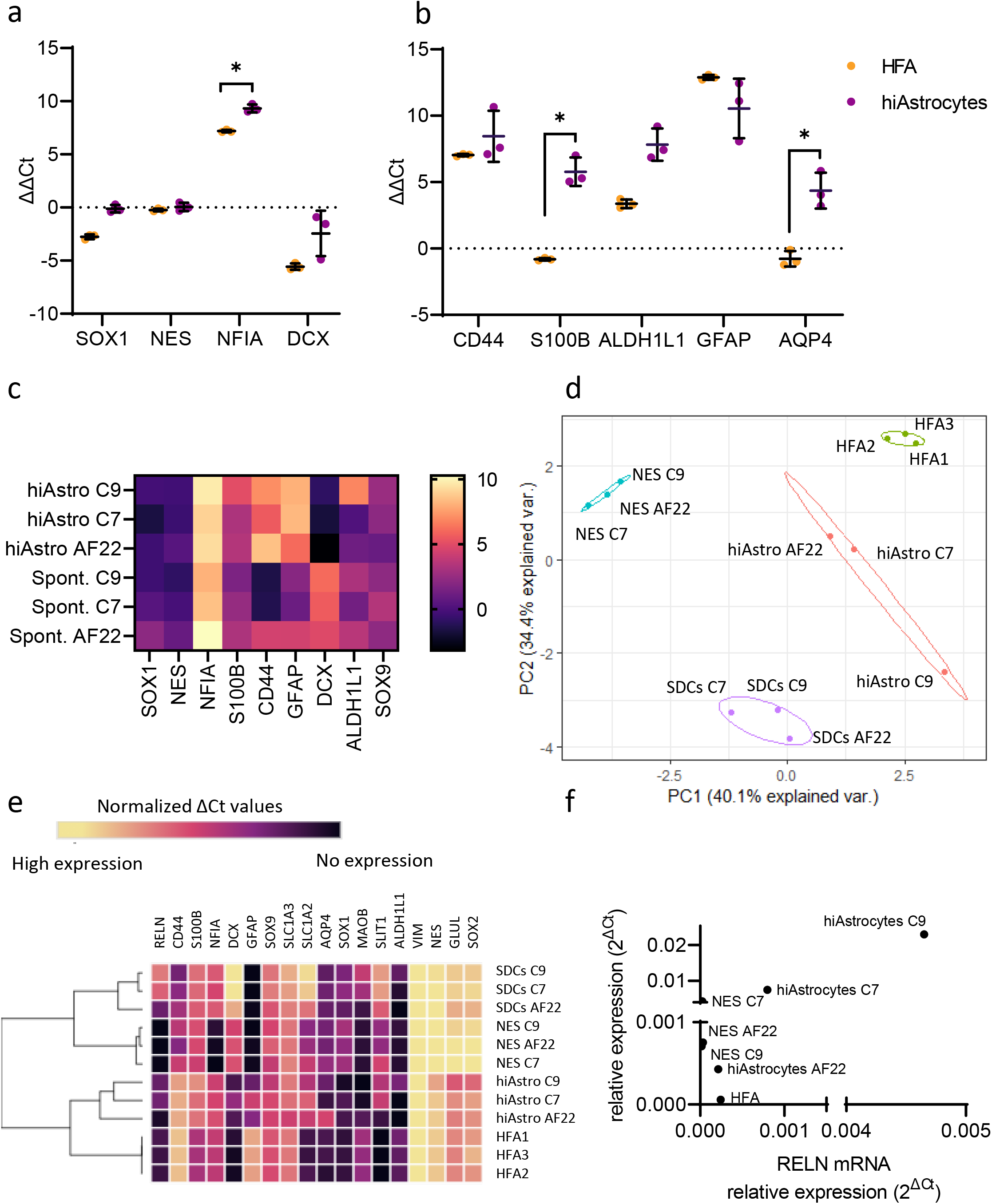
HiAstrocytes have a distinct astrocytic mRNA expression profile that differs from spontaneously differentiated cells. mRNA analysis (ΔΔCt values) of a) *SOX1, NES, NFIA* and *DCX* and b) astrocyte-specific markers *CD44, S100B, ALDH1L1, GFAP and AQP4* for hiAstrocytes C9 (purple) and HFA (orange). Data shown from n = 3 independent experiments. Error bars represent ±SD. Statistical analysis was done on the ΔCt values by using multiple unpaired student’s test (Holm-Šídák method) with Welch correction. *p <0.05. c) Heatmap mRNA levels (ΔΔCt) of hiAstrocytes and SDCs from NES C9, C7 and AF22 d) PCA plot and e) Hierarchical clustering of NES C9, C7 and AF22 along with their corresponding differentiated astrocytes (hiAstro) and spontaneously differentiated cells (SDCs). HFA from one isolation were used. Normalized ΔCt values were used for the PCA plot and hierarchical clustering.

Next, we sought to unveil the transcriptomic differences between the hiAstrocytes, and cells generated by spontaneous differentiation of the NES lines. Spontaneously differentiated cells (SDCs) were generated by growth factor withdrawal for 28 days (Figure S1a). *NFIA* was, strikingly, upregulated in all cell lines in both hiAstrocytes and SDCs, while *CD44* expression was restricted to the hiAstrocytes (all lines) and SDCs AF22 (Figure 2c). As expected, *DCX* was upregulated in all lines in SDCs; conversely, *DCX* was downregulated in hiAstrocytes (all lines). Glial Acidic Fibrillar Protein (GFAP) was highly upregulated in hiAstrocytes (all lines), while *GFAP* expression varied in SDCs. SDCs in AF22 showed higher upregulation of GFAP than SDCs C9 or C7. The differential expression of *GFAP* and *CD44* in SDCs suggests a mixture of neurons and glia in SDCs AF22 while there is a neuronal enrichment in SDCs C9 and C7.PCA of 19 genes analyzed by RT-qPCR showed a distinct separation between SDCs and hiAstrocytes, denoting the impact of the differentiation strategy on enriching astrocytic fate (figure 2d). hiAstrocytes clustered closer to HFA than NES or SDCs. Specifically, hiAstrocytes C7 and AF22 clustered closer to HFA than hiAstrocytes C9. NES and SDCs had distinct clusters further away from HFA. Hierarchical clustering revealed that indeed hiAstrocytes from all three lines clustered with HFA and were distinct from SDCs.

We also assessed the regionality of NES and hiAstrocytes. Gene expression analysis of two regionality markers *RELN* and *SLIT1*, revealed that, strikingly, the NES lines have a different dorsoventral identity (Figure 2e). While all lines had no or insignificant expression of *RELN*, NES C7 seems to have a distinct regional identity denoted by the *SLIT1* expression, which remained almost unchanged during astrocyte differentiation, denoting a regionally patterned precursor that gives rise to specifically VA3 astrocytes. In hiAstrocytes C9, *RELN* and *SLT1* were upregulated, while in hiAstrocytes AF22, there was a very low expression (*RELN* Ct = 33, *SLIT:* Ct = 32). HFA also had a similar RELN expression pattern (Ct = 32), while *SLIT1* was not detected.

### Quantification of astrocytic markers and processes shows a superior proteomic and phenotypic profile compared to HFA

We performed immunocytochemistry and stained hiAstrocytes C9, C7, AF22 and HFA for the common astrocytic markers S100B, CD44, GFAP, AQP4 ALDH1L1 and the cytoskeletal marker VIMENTIN. All hiAstrocytes lines and HFA stained positive for CD44 (Figure 3a) while ICC quantification all models reached almost 100% CD44^+^ cells (Figure 3b). HiAstrocytes C9, C7, AF22 and HFA did not stain for DCX (Figure S1b). While hiAstrocytes and HFA were positive for S100B^+^, hiAstrocytes had a higher percentage of S100B^+^ astrocytes (hiAstrocytes C9/C7/AF22 vs. HFA, 99%/96%/97% vs. 84%). Intensity analysis of each population showed that hiAstrocytes C9 and C7 populations had higher average intensity than HFA, as denoted by the shift in the violin plots (Figure S1c).

**Figure 3.**
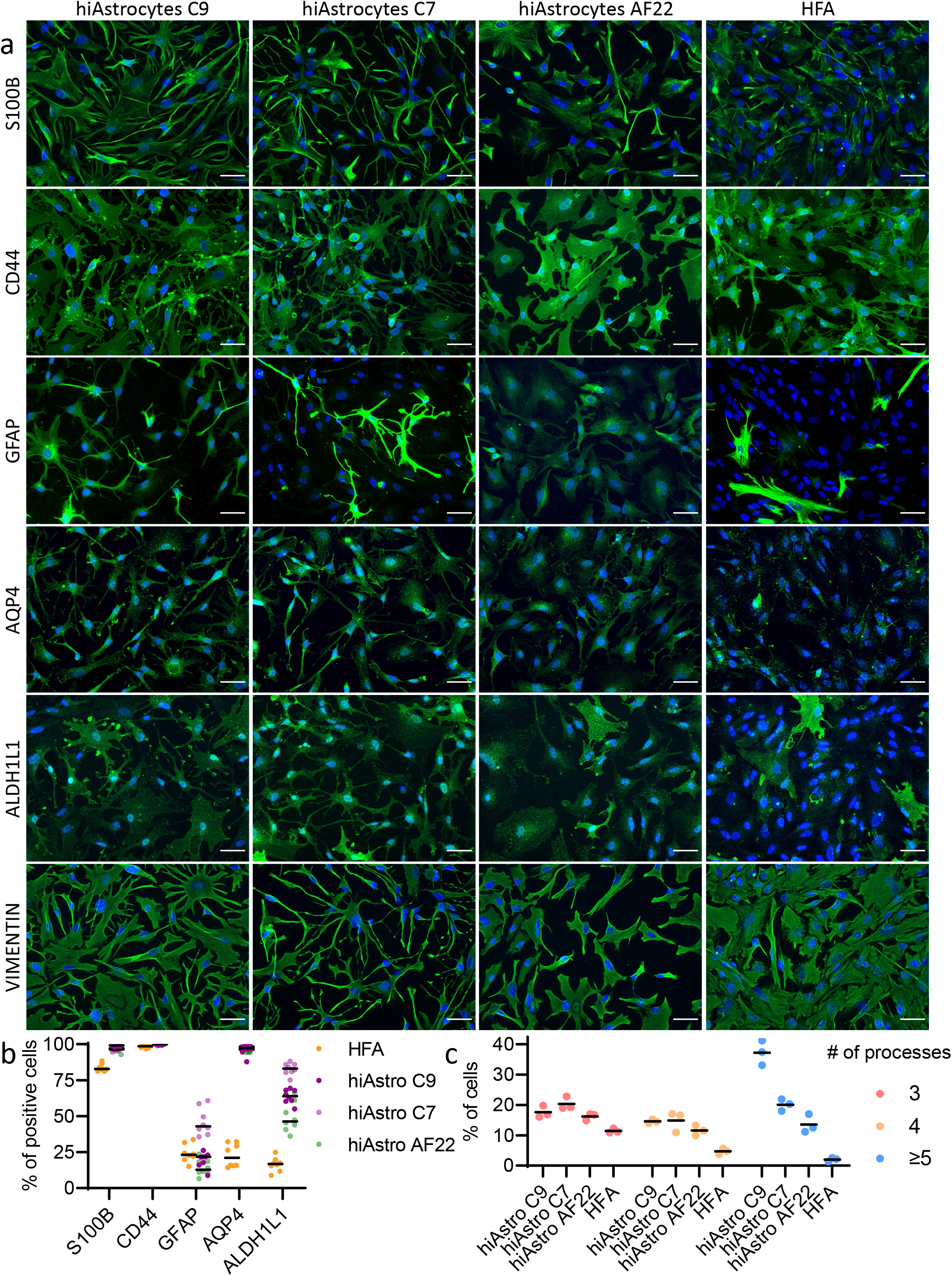
Quantification of astrocytic markers and processes shows superior proteomic and phenotypic profile compared to HFA. a) ICC images of astrocytic markers S100B, CD44, GFAP, ALDH1L1 and the cytoskeletal marker VIMENTIN of hiAstrocytes C9, C7 and AF22 along with HFA. Scale bar, 50 μm. b) quantification of astrocytic markers S100B, CD44, GFAP, AQP4 and ALDH1L1, each dot represents one field of view c) quantification of astrocytic processes, each dot represents the average of four fields of view.

HFA were 24% GFAP^+^ while hiAstrocytes C9, C7 and AF22 were 20%, 46% and 14% GFAP^+^, respectively. Quantification of ALDH1L1 revealed a low percentage (16%) of positive cells while the quantification among the three lines was not coherent, with hiAstrocytes C7 having the higher percentage of positive cells (83%) followed by hiAstrocytes C9 (64%) and AF22 (49%). HiAstrocytes from all three lines were positive for AQP4 with almost 100% positive cells (hiAstrocytes C9/C7/AF22, 96%/98%/96%), while HFA had a lower percentage of AQP4-positive cells (16%).

For the quantification of astrocytic processes, we stained cells with VIMENTIN. HiAstrocytes C9 showed the highest percentage (37%) of multiple processes (>=5, Figure 3c), followed by hiAstrocytes C7 (20%) and AF22 (13%) (>=5, Figure 3d). Only 2% of HFA exhibited multiple processes (>=5, Figure 3d).

#### HiAstrocytes exhibit functional EAAT1- and EAAT2-mediated glutamate uptake

We next sought to characterize the glutamate uptake capacity of hiAstrocytes, a critical in vivo functionality of astrocytes. NES showed an uptake of 7.25 nmol per million cells while hiAstrocytes showed more than a 10-fold increase in glutamate uptake than NES (80.57 nmol per million cells, p<0.0001). HiAstrocytes showed four times higher glutamate uptake rate compared to HFA (20.03 nmol per million cells, p<0.0001)

We used the non-substrate compounds UCPH1 and WAY213613 to inhibit the two astrocytic glutamate transporters EAAT1 and EAAT2, respectively. Both inhibitors significantly blocked glutamate uptake in hiAstrocytes compared to vehicle (vehicle vs. UCPH1, 80.57 vs. 7.13 nmol per million cells, p<0.0001 and vehicle vs. WAY213613, 80.57 vs. 52.95 nmol per million cells, p=0.0057). Even though the GLAST inhibition (UCPH1) lowered glutamate uptake compared to vehicle (vehicle vs. UCPH1, 20.03 vs 5.16 nmol per million cells), it was not significant (p=0.3118). HFA did not exhibit functional EAAT2 with the WAY213613 inhibitor.

Further characterization of those transporters through RT-qPCR and ICC revealed that in hiAstrocytes SLC1A2, the gene encoding for the EAAT2 transporter was highly upregulated (Figure 4b) compared to HFA, which showed a slight downregulation (hiAstrocytes vs. HFA, ΔΔCt 6.46 vs. −1.12). The transporter SLC1A3 had almost the same expression pattern in both cell populations. ICC of the two glutamate transporters showed that both hiAstrocytes and HFA are EAAT1^+^. HiAstrocytes stained positive for EAAT2 while HFA were weakly stained (Figure 4c).

**Figure 4.**
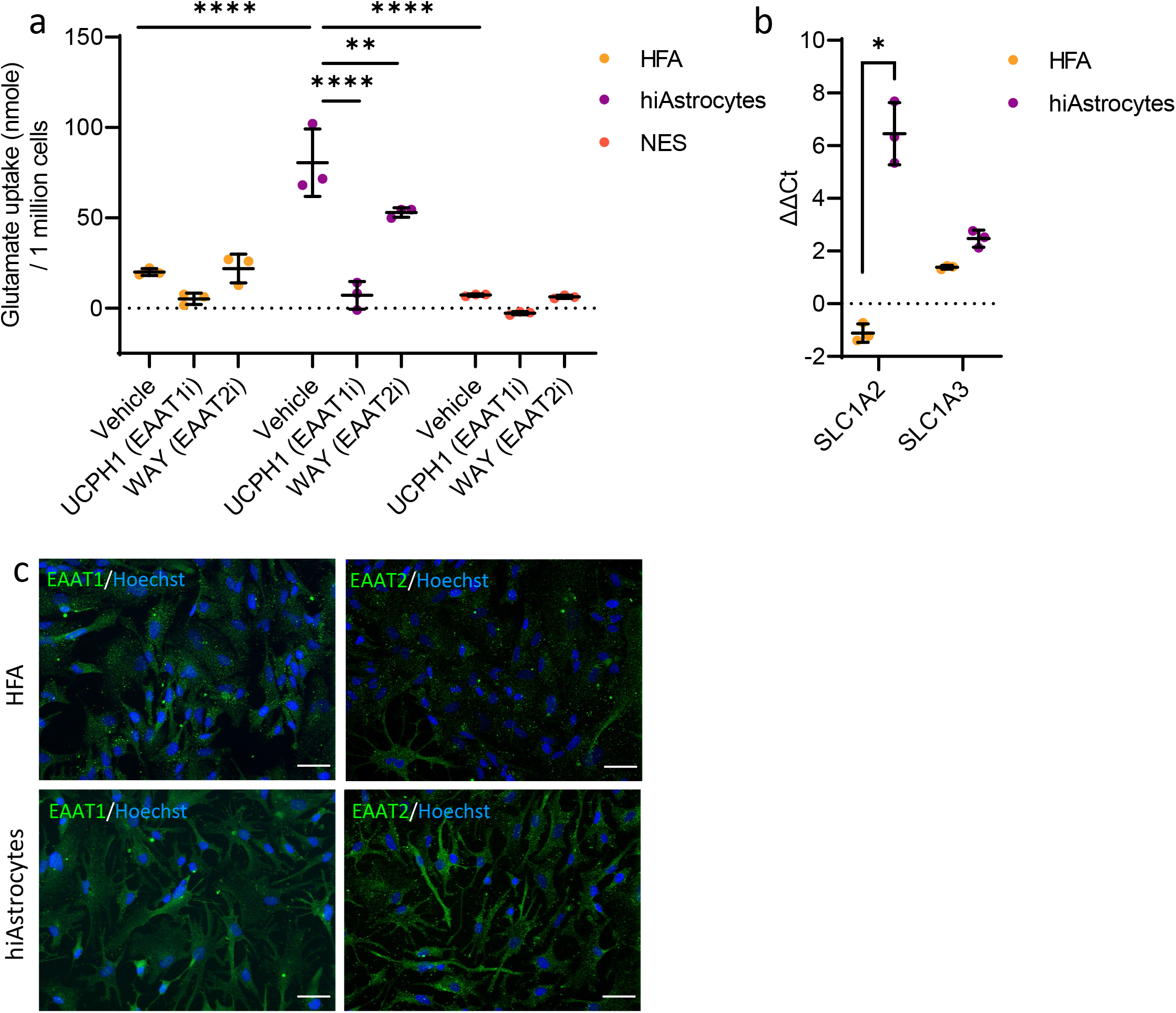
HiAstrocytes exhibit functional EAAT1- and EAAT2-mediated glutamate uptake. a) Glutamate uptake assay for NES C9 (orange), hiAstrocytes C9 (purple) and HFA (yellow), n = 3 independent wells. Error bars represent ±SD. Statistical analysis was done using two-way ANOVA followed by Tukey’s post-hoc test. *p < 0.05, **p < 0.01, ***p < 0.001, ****p <0.0001 b) mRNA analysis of glutamate transporters *SLC1A2* and *SLC1A3* for hiAstrocytes C9 and HFA, n = 3 independent experiments. Error bars represent ±SD. Statistical analysis was done on the ΔCt values by using multiple unpaired student’s test (Holm-Šídák method) with Welch correction. *p <0.05. c) ICC images of glutamate transporters EAAT1 and EAAT2 for HiAstrocytes C9 and HFA.

### HiAstrocytes harbor unique inflammatory potency and antioxidant properties

A vital aspect of a differentiated cell’s repertoire is its capacity to respond to inflammatory agents, which should be on par with the in vivo counterpart, in our case, HFA. That is an important aspect of modeling pathologies in vitro. Hence, we challenged the hiAstrocytes and HFA with IL1-β and stained for the intracellular adhesion molecule 1 (ICAM-1) and measured the cytokine secretion of IL-6 and IL-8, two cytokines that are secreted by astrocytes upon inflammation.

ICC revealed that all lines and HFA showed almost no staining of ICAM-1 when unstimulated, while when incubated with IL1-β for 24 hours, most cells were ICAM^+^ (Figure 5a). Specifically, quantification of the ICC showed that in basal conditions, all hiAstrocytes and HFA had less than 10% ICAM-1^+^(Figure 5b). Upon inflammatory stimuli, hiAstrocytes C9 and C7 had the higher percentage of ICAM-1^+^ cells, 84% and 83%, respectively. HFA had slightly fewer positive cells, amounting to 75% ICAM-1^+^. HiAstrocytes AF22 had the lowest number of ICAM-1^+^,61%.

**Figure 5.**
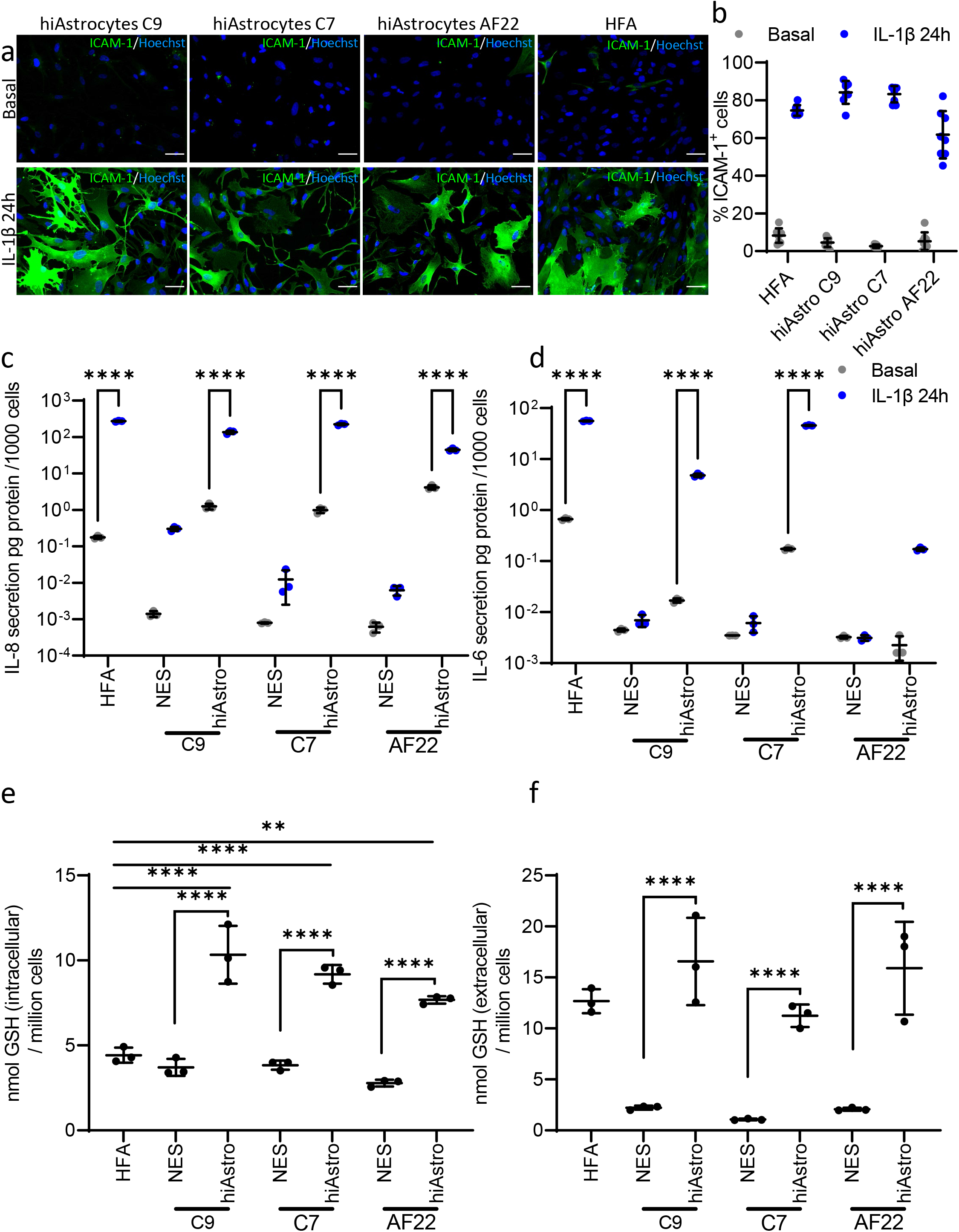
hiAstrocytes harbor unique inflammatory potency and antioxidant properties. a) ICC images of ICAM-1 in hiAstrocytes C9, C7 and AF22 along with HFA under basal and inflammatory conditions (IL-1β 50 ng/ml for 24h). Scale bar 50 μm. b) Quantification of ICAM-1^+^ cells under basal (grey) condition or under inflammatory stimuli (IL-1β 50 ng/ml for 24h, blue). Secretion levels of c) IL-8 and d) IL-6 in NES C9, C7 and AF22 and their corresponding astrocyte differentiation (hiAstrocytes C9, C7 and AF22) and HFA under basal (grey) and inflammatory conditions (IL-1β 50 ng/ml for 24h, blue), n = 3 independent wells. Glutathione levels e) intracellularly and f) extracellularly. NES C9, C7 and AF22 along with their corresponding astrocyte differentiation (hiAstrocytes C9, C7 and AF22) and HFA, n = 3 independent wells. Error bars represent ±SD. Statistical analysis was done using two-way ANOVA followed by Tukey’s post-hoc test. *p < 0.05, **p < 0.01, ***p < 0.001, ****p <0.0001.

Cytokine secretion of IL-8 revealed that hiAstrocytes (all lines) and HFA responded to IL-1β by secreting IL8, which was significant compared to basal conditions (p>0.0001, figure 5c). All lines showed a differentiation-acquired immune potency of IL-8 secretion. HFA and hiAstrocytes C7 had the highest secretion (56 pg and 47 pg per 1000 cells, respectively) of IL-6 (Figure 5d). HiAstrocytes C9 secreted an order of magnitude less, amounting to 8 pg per 1000 cells, while hiAstrocytes AF22 secreted 0.171 pg per 1000 cells. IL-1β-challenged HFA, hiAstrocytes C9 and C7 had significantly higher IL-6 secretion compared to their basal conditions (p>0.0001), while IL-6 secretion in hiAstrocytes AF22 challenged with IL-1β was not significantly higher compared to their basal conditions (p>0.001, Basal: 0.013 vs. 0.171 pg per 1000 cells).

A largely unexplored astrocytic aspect in stem cell-derived astrocytes is the capacity of astrocytes in vivo to protect the CNS against oxidative stress through glutathione. Astrocytes have a very high glutathione content (~8mM) (Dringen and Hamprecht, 1998) that they continuously replenish since they constantly provide GSH precursors to neurons and brain endothelial cells(Huang *et al*., 2020).

We measured the total GSH content of NES, hiAstrocytes and HFA, and we also sampled medium extracellularly to assess 1) the intracellular content of hiAstrocytes and how it compared to HFA GSH levels and 2) whether they can export it and thus extend their capacity to protect neighboring cells against reactive oxygen species.

Our analysis showed that all hiAstrocytes showed significantly higher intracellular total GSH than the undifferentiated NES (p<0.0001, Figure 5e). When compared to HFA, hiAstrocytes had higher total GSH intracellularly, while HFA had 4.42 nmol total GSH per million cells, hiAstrocytes C9, C7 and AF22 showed 10.33 (p<0.0001), 9.18 (p<0.0001) and 7.68 nmol (p=0.0014) total GSH per million cells, respectively.

Measurements of total extracellular GSH revealed that all hiAstrocytes exported considerably more total GSH than their NES counterparts. While all NES lines hovered between 1.1-2.2 nmol total GSH per million cells, hiAstrocytes C9, C7 and AF22 showed a remarkable increase ranging from 11.2 to 16.6 (p<0.0001, figure 5f). Expanding more on that, the total extracellular GSH in hiAstrocytes C9 and AF22 was almost eight times higher than their respective NES lines. HiAstrocytes C7 showed an increase amounting to 10 times more compared to NES C7. Extracellular values of hiAstrocytes were on par with HFA.

## Discussion

Over the past years, there has been a plethora of astrocytic differentiation protocols, some more elaborate and time-consuming than others, spanning over many months or sorting steps(Holmqvist *et al*., 2015; Santos *et al*., 2017; Leventoux *et al*., 2020). Our differentiation strategy relies on creating an astrocytogenic milieu that brings forth astrocytic traits in only 28 days. The differentiation protocol we report is advantageous for CNS disease modeling, considering the simplicity and relatively cost-effective reagents. We created the astrocytogenic milieu as a combination of ECM component, culture medium and cell-to-cell communication (regulated via seeding density) to generate functional and morphologically star-shaped astrocytes in under 28 days. Collagen does not promote neuronal differentiations and is secreted by astrocytes in vitro. Gelatin, combined with a growth media designed to promote and sustain astrocytes creates an environment that guides NES into functional star-shaped astrocytes.

In our previous work, we compared astrocyte differentiation protocols to adult astrocytes per industry standards(Lundin *et al*., 2018). However, fetal astrocytes are a more relevant model to compare hiPS-derived astrocytes. Adult human astrocytes have undergone a developmental process that is very challenging to recapitulate *in vitro*; hence, astrocytic differentiations are most likely to generate astrocytes that have developmental proximity to human fetal astrocytes than to adult human astrocytes.

Our differentiation strategy generates astrocytes that resemble HFA both on an mRNA and protein level. HiAstrocytes and HFA are not in the same cluster In the PCA plot, albeit hierarchical clustering reveals that hiAstrocytes cluster with HFA and not with SDCs. We postulate that the reasons behind that deviation are: 1) the different developmental stage, 2) different regionality, and 3) the fact that there is always a caveat with the acute isolation of primary cells (Lange *et al*., 2012). Additionally, *in vitro* expansion of primary astrocytes follows an adaptation mechanism that may ensue a transcriptomic shift (i.e. dedifferentiation) similar to what has been seen with other CNS primary cells such as brain endothelial cells (Sabbagh *et al*., 2018).

HiAstrocytes had a higher expression of *S100B* compared to HFA. The S100B enrichment in hiAstrocytes corroborates with our ICC data where there is a higher percentage of S100B^+^ cells than HFA. Interestingly, hiAstrocytes had a higher average intensity of S100B^+^ cells (per cell) than HFA. That could denote a developmental stage that surpasses the developmental stage of commercially available HFA (18-20 GW). In mice, *S100B* expression of GFAP^+^ astrocytes is associated with a mature state that lacks neural stem cell traits (Raponi *et al*., 2007). In humans, the transition from fetal to mature astrocytes is characterized by S100B upregulation (Zhang *et al*., 2016), while the higher percentage of AQP4^+^ and ALDH1L1^+^ in hiAstrocytes reinforces the notion that hiAstrocytes have a developmental stage that transcends the 20 GW developmental stage of HFA (Zhang *et al*., 2016).

Secondly, a largely unaddressed issue in most hiPS-derived astrocyte protocols is the regionality of astrocytes. Transcriptomic analysis of two regionality markers associated with the spinal cord, *RELN* and *SLIT1* (Hochstim *et al*., 2008; Rowitch and Kriegstein, 2010; Clarke *et al*., 2021), revealed that the hiAstrocytes have a distinct regional identity; specifically, hiAstrocytes C9 appeared to be either a mixture of VA1, VA2 astrocytes or VA3. Interestingly, NES C7 was the only NES line that showed expression of *SLIT1* which persisted during the differentiation associating hiAstrocytes C7 with ventrally located VA3 population. AF22 HiAstrocytes did not show expression of either *RELN* or *SLIT1*, suggesting a more anterior regional identity. HFA did not express any of these markers, as expected since these astrocytes were isolated from the cortex. However, more regionality markers should be explored to pinpoint the regionality of hiAstrocytes accurately. To what extent neural stem cells patterning affects the anteroposterior and dorsoventral identity of hiAstrocytes remains to be seen. Bradly and et al. showed that the regionality of neural stem cells influences the gene expression of downstream differentiation to astrocytes; hence, it follows that astrocytes derived from differentially patterned NES lines cannot possibly cluster very tightly (Bradley *et al*., 2019) owing to differential expression patterns in various regions of the brain, one notable example is glutamate transporter expression(Bar-Peled *et al*., 1997) which is region-dependent.

The non-overlapping clusters in the PCA echo the abovementioned differences between hiAstrocytes and HFA; perhaps inclusion of primary astrocytes of different parts of the brain would better associate with hiAstrocytes. Albeit having a different regional identity, all hiAstrocytes stained positive for the canonical astrocytic markers, S100B, AQP4, ALDH1L1 and CD44. Interestingly, the generated cells are GFAP^+^ which was not observed in our previous protocol to generate astrocytes from long-term proliferating NES(Lundin *et al*., 2018, 2020; Lam, Sanosaka, *et al*., 2019). NES (all lines) did not stain for any of these markers but were positive for VIMENTIN (Figure S2).

The morphology of astrocytes is also another salient feature of astrocytes, mirroring their physiological/pathological state. A star-shaped morphology has been greatly elusive in shorter differentiation protocols (Tcw *et al*., 2017; Lundin *et al*., 2018), and only possible to attain after prolonged differentiation ~5 months as documented by Oksanen and et al.(Oksanen *et al*., 2017). The hiAstrocytes showed a star-like morphology and had more complex morphologies (processes >= 3, 41 - 69%, line depended, Figure S3) than HFA (18%), making this differentiation strategy ideal for CNS disease modeling where astrocytic processes are affected by pathological conditions(Oberheim *et al*., 2008; Rodríguez *et al*., 2009; Schiweck, Eickholt and Murk, 2018). A noteworthy pathological condition is schizophrenia, where researchers have shown that astrocytes in schizophrenia have a lower number of astrocytic processes than healthy astrocytes (Windrem *et al*., 2017).

A noticeable difference between the NES lines in SDCs was the different populations they generated. NES C9 and C7 were biased towards neuronal commitment, while NES AF22 generated a mixture of glial and neuronal populations. Our data corroborate with findings from other studies; specifically, NES AF22 has been shown to produce a mixture of glial and neuronal population upon growth factor withdrawal(Lam, Sanosaka, *et al*., 2019) while NES C7 mainly generated neurons(Lam, Moslem, *et al*., 2019). Adhering to that paradigm, NES AF22 should have the greatest potential for astrocyte differentiation (compared to NES C7 and NES C9), but strikingly NES AF22 underperformed compared to NES C9 and NES C7. NES AF22 has previously also underperformed compared to other lines when differentiated towards the astrocytic lineage(Lundin *et al*., 2018).We postulate that these differences could be attributed either to the different neural inductions used to generate the lines (NES C9 and C7: dual-SMAD and NES AF22: spontaneous neural induction) or to the type of reprogramming of the corresponding hiPSC lines (iPS C9 (Uhlin *et al*., 2017) and C7 (Kele *et al*., 2016) non-integrating sendai virus and iPS AF22 integrating lentivirus). The former could be a plausible reason since differences in neural inductions has been shown to affect astrocytic potential (Nadadhur *et al*., 2018); however, more experiments on how neural induction/reprogramming affects downstream differentiations should be done.

EAAT1 expression and functionality has been documented previously in NES-derived astrocytes (Lundin *et al*., 2018) and constitutes perhaps the bare minimum functional requirement of iPS-derived astrocytes. HiAstrocytes showed specific EAAT1 glutamate uptake and had a similar expression pattern to HFA. EAAT2 is not expressed in HFA *in vitro* (Lee *et al*., 2008), and its expression is induced by coculturing primary astrocytes with neurons (Swanson *et al*., 1997; Schlag *et al*., 1998) or brain endothelial cells (Lee *et al*., 2017). Interestingly, our results show that EAAT2 was significantly upregulated in hiAstrocytes than HFA, without using any molecular inducers such as Ceftriaxone (Lee *et al*., 2008) or coculturing with other CNS cells. The glutamate assay and the specific EAAT2 inhibitor, WAY213613, revealed that EAAT2 transporters in hiAstrocytes are functional. We postulate that the differentiation strategy, more specifically, the high density that cells are kept before each passage and the presence of bFGF in the differentiation media, work synergistically towards EAAT2 expression and functionality. Notch signaling has been shown to increase GLT-1 expression in mice (Lee *et al*., 2017), and the same has been shown for bFGF (Savchenko *et al*., 2019). Other groups have shown EAAT2 expression in iPS-derived astrocytes (Roybon *et al*., 2013; Shaltouki *et al*., 2013; Leventoux *et al*., 2020), and here we report on an astrocytic model that shows specific EAAT2 expression and functionality.

EAAT2 has gained much attention since studies suggest that excitotoxicity-induced neuronal death is closely correlated with neurological disorders (Pajarillo *et al*., 2019) such as ALS and AD (Rothstein *et al*., 1995; Takahashi *et al*., 2015; Garcia-Esparcia *et al*., 2018). In autism, the significance of EAAT2 over EAAT1 has been documented in a mouse model (Bristot Silvestrin *et al*., 2013). EAAT2 expression and functionality is pivotal when attempting to model in vitro these neurological disorders. Ceftriaxone is one candidate that showed promising results in vitro and animal models in increasing GLT-1 expression (Rothstein *et al*., 2005; Colton *et al*., 2010). Even though this specific compound failed in clinical trials, compounds that modulate EAAT2 expression are still a plausible therapeutic route for ALS (Rosenblum and Trotti, 2017, p. 2).

Additionally, hiAstrocytes exhibited strong inflammatory potency. Upon IL-1b simulation, hiAstrocytes secreted IL-6 and IL-8, which is on par with what other groups have shown for hiPS-derived astrocytes (Holmqvist *et al*., 2015; Santos *et al*., 2017; Perriot *et al*., 2018). We quantitatively analyzed and compared cytokine secretion between hiPS-derived astrocytes and primary fetal astrocytes. Our results showed that hiAstrocytes C9 and C7 secreted IL-8 and IL-6 in comparable levels to HFA. Additionally, upon IL-1b stimulation, hiAstrocytes expressed ICAM-1. ICAM-1 upregulation is a link in the relay of inflammatory stimuli between contact-mediated and secreted cytokines, activation of ICAM-1 has been shown to elicit IL-6 secretion (Lee *et al*., 2000). Additionally, NES (all lines) were not responsive when challenged with IL-1β staining negative for ICAM-1.

The intracellular GSH content of hiAstrocytes and HFA is on par with other astrocytes studies (Figure S3a), ranging from 16 to 50 nmol/mg protein (Raps *et al*., 1989; Devesa *et al*., 1993; Dringen, 2000). HiAstrocytes C9 and C7 hovered around the higher end of that range (57 nmol/mg protein), while hiAstrocytes AF22 and HFA had lower GSH content,35, and 24 nmol/mg protein, respectively. Our differentiation approach generates astrocytes that exhibit superior export of GSH. One other study that measured extracellular GSH in hiAstrocytes documented ~8.13 nmol/mg protein (Oksanen *et al*., 2017) while our study generated astrocytes that exported at least six times more GSH (Figure S3b and c). The rate of GSH export is hiAstrocytes ranged between 2.9 to 3.8 nmol GSH/ (h x mg protein) (Figure S3d). HFA showed an export rate of 2.9 nmol GSH / (h x mg protein).

Interestingly, our values are in the range of previous studies in rat primary astrocytes, 3.2 nmol / (h x mg protein) (Dringen, Kranich and Hamprecht, 1997). HiAstrocytes C9 and C7 exhibited ~7 times more GSH export, than their respective NES lines and AF22 ~10 times more GSH than NES AF22 (in nmol GSH / million cells). This data could suggest that differentiated cells have a proper expression of *ABCC1* for GSH export (Renes *et al*., 1999; Hirrlinger, Schulz and Dringen, 2002; Minich *et al*., 2006). Astrocytes continuously synthesize GSH; the highest consumption of GSH occurs in the form of GSH export. Hence astrocytes synthesize steady-state GSH to compensate for exported GSH. The intracellular content of GSH is defined by 1) the rate that cells synthesize GSH from precursor molecules and 2) the rate that cells export GSH. Contemplating the similar export rates between hiAstrocytes and HFA and considering that all cells were cultured in the same media (i.e., same availability of GSH precursors), hiAstrocytes showed 2 to 3 times more GSH content in 24 hours, compared to HFA. This difference suggests that HFA lag and cannot fully compensate for the GSH export; hence hiAstrocytes have superior capacity to mitigate insults and reactive oxygen species. The strategic location of astrocytes in the brain, covering over 90% of the brain vessels with their processes, constitutes astrocytes the first line of defense against xenobiotics and toxins that enter the brain. Neurons are unable to synthesize glutathione on their own and rely on astrocytes as a glutathione source (Wang and Cynader, 2000). Moreover, astrocyte-secreted glutathione counteracts the detrimental effects of an insult to the blood-brain barrier (Huang *et al*., 2020). ROS are prevalent in many, if not all, neurological conditions (Bains and Shaw, 1997; Cadet and Brannock, 1998; Li *et al*., 2013; Popa-Wagner *et al*., 2013; Fang *et al*., 2017), are they part of the etiology or disease progression? Are astrocytes the cause or the domino factor in disease progression? GSH cycle disturbances can potentially shed light on this front.

We report on specific astrocytic traits that have not been assembled before in a hiPS-derived astrocyte generation. Astrocytes harbor in their immune response a repertoire that transcends mere cytokine secretion; astrocytic responses are characterized by a cascade of reactions that involve morphological rearrangement of their processes and expression of adhesion molecules (e.g., ICAM-1), loss of glutamate transporters and glutathione redox balance shift. Consequently, this hiPS-derived astrocytic protocol constitutes a multifaceted *in vitro* model that may serve as a powerful tool enabling pathological cues to surface and potentially further deepen our knowledge of how astrocytes are involved in the etiology, onset, and progression of CNS pathological conditions. This conveniently short differentiation protocol may contribute to the advancement of personalized medicine and to improved implementations of more complex vitro models such as organ-on-chip technologies.

## Acknowledgements

We thank the iPS core and the Biomedicum Imaging Core (BIC) at Karolinska Institute for access to their service.

We also thank Thomas E. Winkler for sharing his knowledge in statistics and Isabelle Matthiesen for the help with cell culture.

PN and AH acknowledges funding from the Swedish Research Council (2019-01803).

AH acknowledges funding from the Knut and Alice Wallenberg Foundation (no. 2015-0178), Göran Gustafsson Stiftelse, and Forksa utan Djurförsök.

## Author contributions

DV: Conceptualization; methodology; investigation; analysis and visualization; writing – original draft.

PN: Validation; methodology; writing review & editing

AH: Conceptualization; methodology; resources, writing review & editing, Supervision and Administration.

## Figure legends

**Figure S1.**
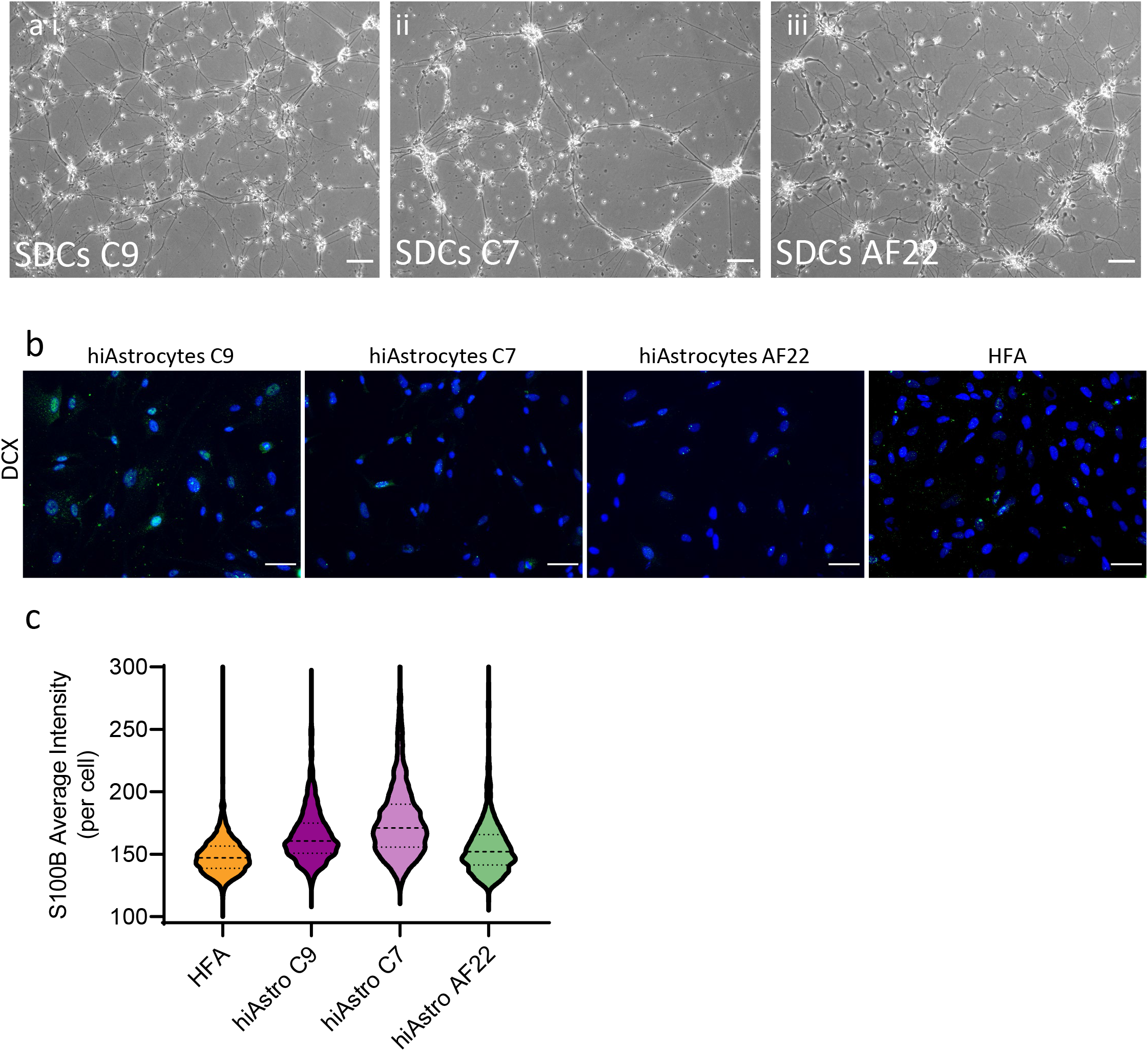
Characterization of SDCs and hiAstrocytes. a) Brightfield images after 28 of spontaneous differentiation of i) NES C9 (SDCs C9) ii) NES C7 (SDCs C7) and iii) NES AF22 (SDCs AF22). Scale bar 100 μm. b) DCX staining of hiAstrocytes C9, C7, AF22 and HFA. Scale bar 50 μm. c) Violin plot depicting the average intensity (per cell) of S100B for hiAstrocytes C9, C7, AF22 and HFA.

**Figure S2.**
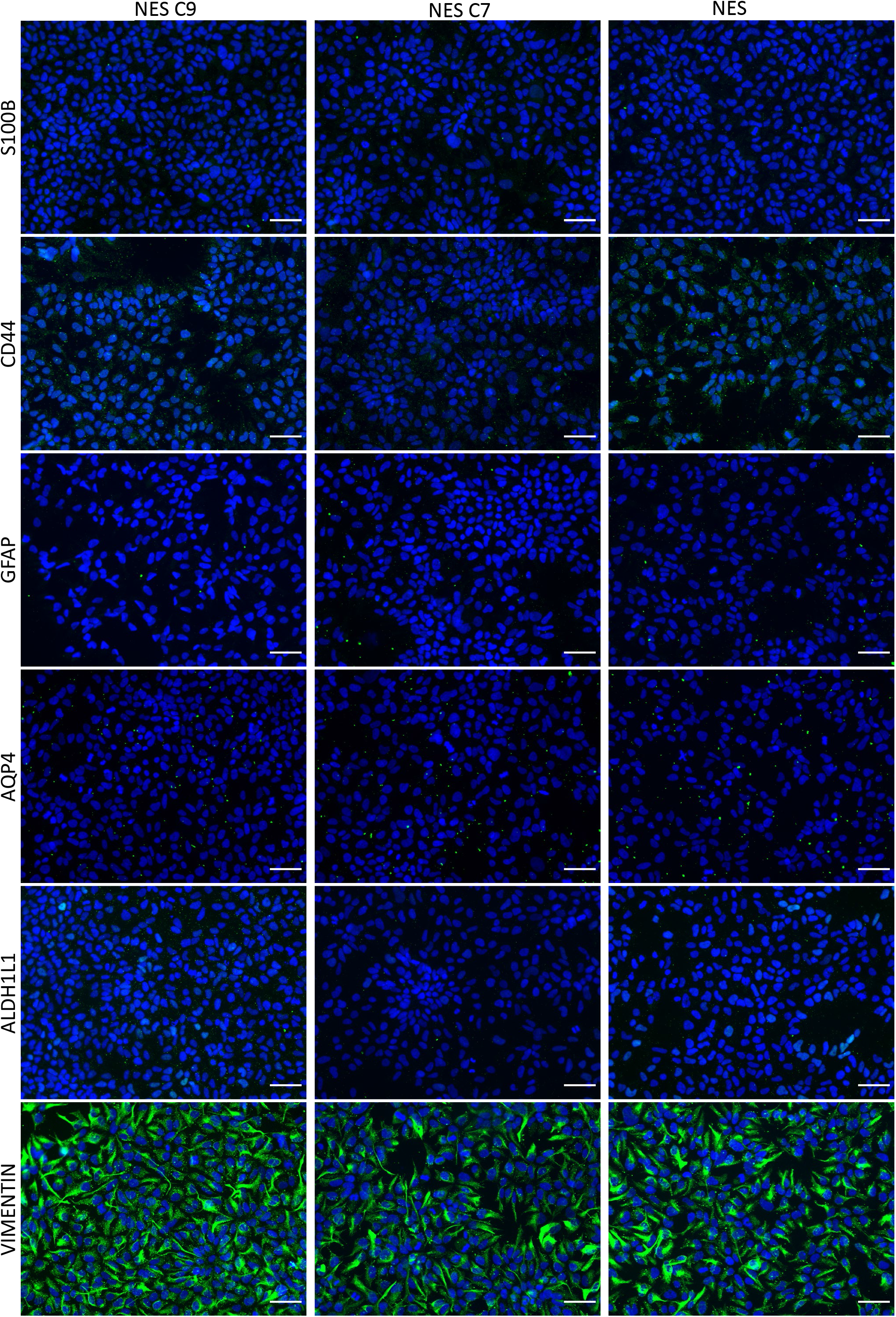
ICC of the NES lines used in this study. ICC images of NES lines NES C9, C7 and AF22 for the astrocytic markers S100B, CD44, GFAP, ALDH1L1. Scale bar 50 μm (20x).

**Figure S3.**
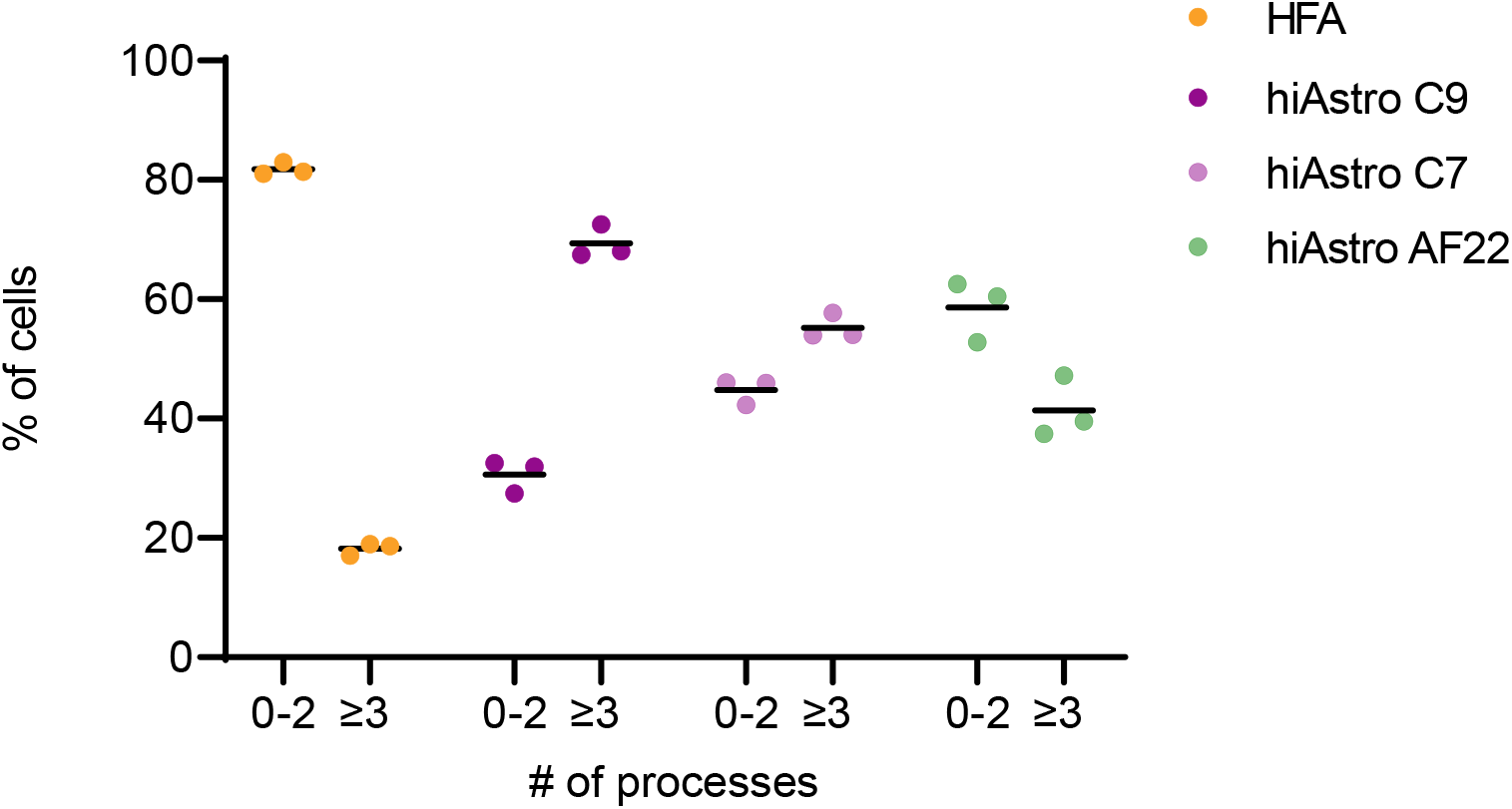
Quantification of astrocytic processes. Quantification of astrocytic processes of HiAstrocytes C9 (dark purple), C7 (light purple), AF22 (light purple) and HFA (green). Each dot represents an average of four fields of view (10x).

**Figure S4.**
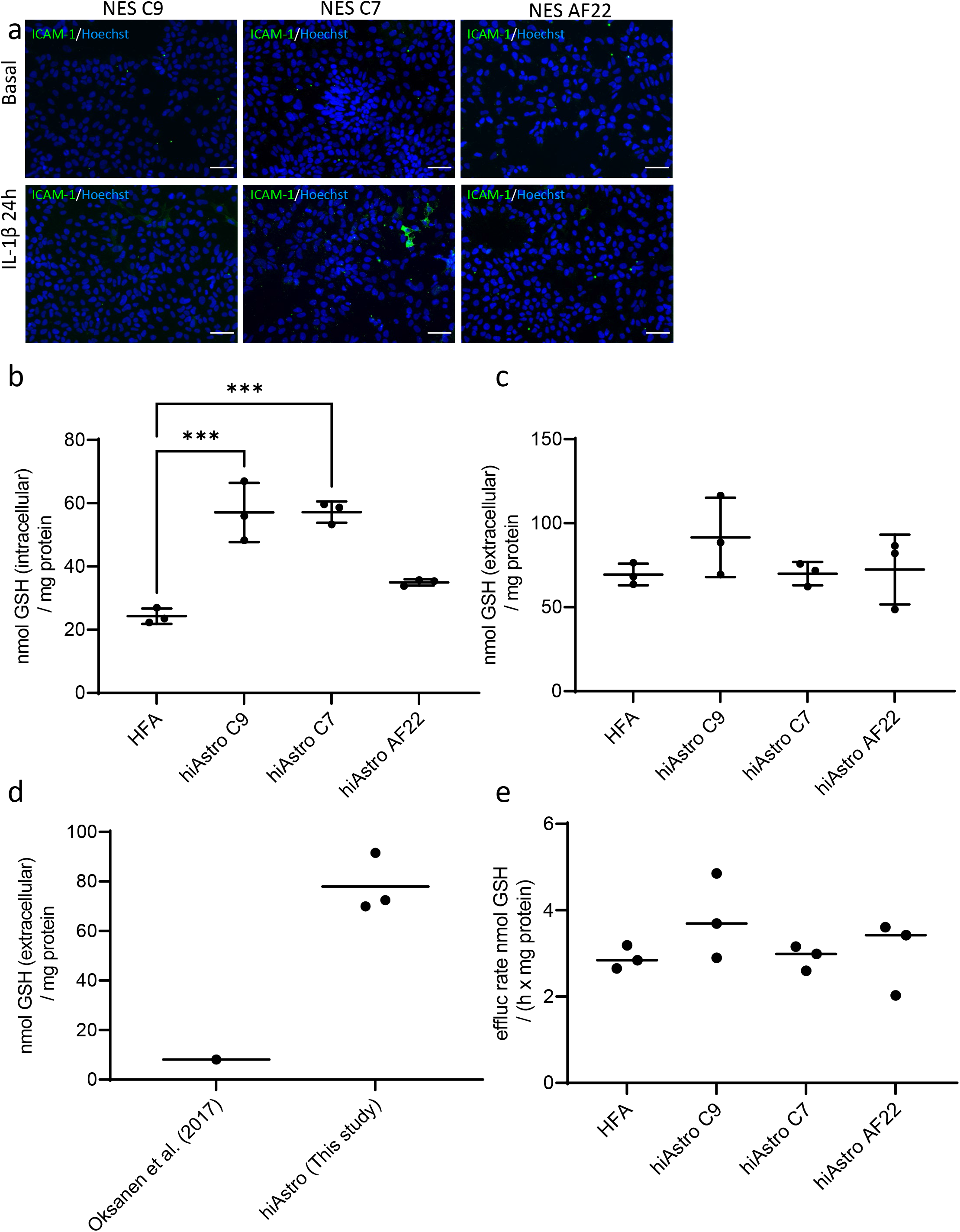
Characterization of immune potency of NES and quantification of glutathione levels of hiAstrocytes and HFA. a) ICAM staining for basal and inflammatory conditions for NES C9, C7 and AF22. Scale bar 50 μm. b) total intracellular and c) total extracellular glutathione content for hiAstrocytes C9, C7, AF22 and HFA normalized by total protein content, n=3 independent wells. Error bars represent ±SD. Statistical analysis was done using two-way ANOVA followed by Tukey’s post-hoc test. *p < 0.05, **p < 0.01, ***p < 0.001, ****p <0.0001. d) comparison with the work from Oksanen et al. (2017), n = 3 different lines, the dot in the Oksanen study is the mean of the values in the Oksanen study derived by dividing with the molecular weight of glutathione e) Glutathione efflux rate of hiAstrocytes C9, C7, AF22 and HFA.

## STAR methods

All reagents were purchased from Thermo Fisher Scientific, MA, USA, unless otherwise stated.

All processes were done according to the manufacturer’s instructions, unless otherwise stated.

### Cell culture

#### Human Neuroepithelial stem cells (NES)

NES lines (Control 9 and 7: male, dual-SMAD neural induction (Chambers *et al*., 2009) and AF22: female, spontaneous neural induction (Falk *et al*., 2012)) were provided by the iPS Core Facility (Karolinska Institute). All lines were cultured and passaged until p.23-24 in DMEM: F12 Glutamax supplemented with N2 1:100, B27 1:1000, 10 ng/ml bFGF (R&D Systems, MN, USA) and 10 ng/ml EGF (Sigma Aldrich, MO, USA) the complete media is termed N2B27 on double coated PLO (20ug/ml Sigma Aldrich, MO, USA) and L2020 (1:500, Sigma Aldrich, MO, USA) flasks. Culture vessels were incubated for 2 hours with PLO washed thoroughly x2 with DPBS (w/o Ca^++^ and Mg^++^) and then coated for at least 2 hours with L2020. After p.23-24, NES were cultured in N2B27 in low EGF (1ng/ml).

NES were passaged 1:4-1:5. Briefly, cells were washed with DPBS (w/o Ca^++^ and Mg^++^) and then incubated with trypLE for 3-4 min. TrypLE was deactivated using equal volumes of DTI and DMEM: F12 glutamax (1:1:1), spun down at 200g and resuspended in N2B27.

The day after passaging bFGF (10ng/ml) or EGF (1 or 10 ng/ml, depending on the passage number) was added cultures the day after the media was completely replenished, this motif of cell feeding was done throughout the NES culture.

#### Astrocyte differentiation

NES were passaged using trypLE for 3-4 min and trypLE was deactivated by adding DTI. Cells were spun down and resuspended in N2B27 with low EGF (1ng/ml). Cells were seeded out in 6-well plates at 288K/well. The day after, cells were carefully washed with DPBS (w/ Ca^++^ and Mg^++^) and media was changed to AM medium (with the addition of AGS and FBS, ScienCell, CA, USA). Media changes were done every other day, and cells were passaged 1-2 days upon reaching confluency, except for the first passage where cells were passaged on days 6-7. For all passages, cells were consistently seeded out at 30K/cm^2^.

All astrocyte differentiations were carried out using NES with p.# >30.

#### Spontaneously differentiated cells (SDCs)

NES (p.#> 30) were detached using trypLE for 3-4 mins and were seeded out at 30K/cm^2^ in double-coated PLO (20ug/ml) and L2020 (1:100) in N2B27 (low EGF). The day after, media was changed to DMEM: F12 glutamax with N2 1:100 and B27 1:100. The media was completely changed every other day. Cells were split once during the differentiation (day 7) at 45K/cm^2^ on PLO-L2020 plates. Following the first passage, media was changed every 3 to 4 days. DMEM: F12 glutamax with N2 1:100 and B27 1:100 was first warmed at 37°C, and then 10ng/ml BDNF and 10ng/ml GDNF (both R&D Systems, MN, USA) was added to the media prior to complete media change. On day 28, cells were harvested.

#### Human Fetal Astrocytes (HFA)

Human astrocytes were cultured in ScienCell Media AM supplemented with 2% FBS, 1% AGS and 1% P/S. HFA that exceeded passage 7 were not used for experiments. Cells were passaged once they reached confluency with a 1:4 ratio. Briefly, cells were washed with DPBS (w/o Ca^++^ and Mg^++^) and then incubated with trypLE for 3-4 min. TrypLE was deactivated using DTI 1:1 (TrypLE:DTI), spun down at 200g and resuspended in complete AM media.

### Immunocytochemistry

Cells were initially washed with DPBS (w/ Ca^++^ and Mg^++^) and fixed with 4% PFA (VWR, PA, USA) for 10 min at room temperature. After washing twice with PDBS (w/ Ca^++^ and Mg^++^), cells were incubated with blocking buffer (10% goat serum (Sigma Aldrich, MO, USA) and 0.1% Triton X-100 in DPBS (w/ Ca^++^ and Mg^++^). Primary antibody incubation was done in dilution buffer (10% blocking buffer), overnight at 4°C. Secondary antibody (Sigma Aldrich, MO, USA) incubation was done in dilution buffer at room temperature for an hour. After 2x washes in DPBS (w/ Ca^++^ and Mg^++^) cells were stained with Hoechst (1:2000) in dilution buffer. Cells were washed 3x and imaged with ImageXpress Micro (Molecular Devices, CA, USA). Quantification of astrocytic markers and astrocytic processes was done by using the Multi-Wavelength Cell Scoring and Neurite Outgrowth module, respectively. A detailed list of the antibodies used can be found in Table S1.

### mRNA expression analysis

Cells were collected, lysed and total RNA was extracted using the RNeasy Mini kit (Qiagen, Germany); cDNA synthesis was carried out using the High-capacity RNA-to-cDNA kit on a thermal cycler (VWR, PA, USA). cDNA samples were analyzed using MySpec (VWR, PA, USA). TaqMan probes of interest were incubated with cDNA samples in Fast Advanced Master Mix. For all samples, GAPDH TaqMan probes were included as reference. Samples were run on a BioRad CFX96 Touch Real-Time PCR Detection system using the multiplex option for superior Ct quantification. Depending on the graph presented, either ΔCt or ΔΔCts were calculated. Samples that lacked Ct values (e.g., *RELN, GFAP* for all NES lines) or had values over 35 were assigned a Ct value of 35 for the ΔΔCt analysis to be possible (or to minimize overestimation of results). PCA was done with R using the prcomp function (R Core Team (2021). R: A language and environment for statistical computing. R Foundation for Statistical Computing, Vienna, Austria. URL https://www.R-project.org/) and hierarchical clustering with the Morpheus software using average linkage clustering, person correlation (Morpheus, https://software.broadinstitute.org/morpheus). A list of the TaqMan probes used can be found in Table S2.

### Glutamate uptake

NES C9, hiAstrocytes C9 and HFA were seeded out in 96-well plates in their respective media. After 72h cells were washed once with HBSS (w/ Ca^++^ and Mg^++^) and then incubated in either vehicle (DMSO) or inhibitors UCPH1 (1uM, EAAT1 inhibitor, Abcam, UK) and WAY 213613 (1.5uM, EAAT2 inhibitor, R&D Systems, MN, USA) for 30 min in HBSS (w/ Ca^++^ and Mg^++^). After that, HBSS was changed with HBSS (w/ Ca^++^ and Mg^++^) containing 50uM of glutamic acid with either vehicle (DMSO) or inhibitors and incubated for 60 min. Samples were taken and analyzed with the glutamate assay kit (Abcam, UK). Following sample collection, cells were incubated with Image-IT™DEAD Green™. Cells were consequently fixed and stained with Hoechst. Images were captured within 24-48h with ImageXpress Micro 10x (Molecular Devices, CA, USA). Live dead count was assessed by the Live/Dead module.

### Inflammatory assay

NES, hiAstrocytes and HFA were seeded out in 96-well plates, and 48h after cells were washed once with DPBS (w/ Ca^++^ and Mg^++^) and challenged with IL-1β (50 ng/ml, R&D Systems, MN, USA). For the basal conditions, cells were washed with DPBS (w/ Ca^++^ and Mg^++^), and media was replenished. After 24h, samples were collected and snap-frozen. Samples were analysed using the Mesoscale system (U-PLEX plate) according to kit instructions. The condition termed “IL-1β 24h” for analysis of ICAM-1 also followed the same procedure. Cell numbers were determined in the same procedure as in the glutamate assay.

### Glutathione assay

NES, hiAstrocytes and HFA were seeded out in T12.5 flasks in their respective media for 48h; cells were washed with (w/ Ca++ and Mg++) all cells were changed to N2B27 with low EGF (1ng/ml) and B27 without antioxidants, control media was also included in a cell-free T12.5 flask. After 24h, media samples were collected from all cells and control media, and cells were harvested and snap-frozen. Media samples and cell suspensions were analyzed kinetically with the Glutathione assay kit (Sigma Aldrich, MO, USA) according to the manufacturer’s instructions with the following modifications: the kinetic reactions were analyzed for 30 min and the calibration curve was adjusted to include more points. For extracellular quantification, the cell-free media flask values were subtracted from the media samples. Deproteinized samples were reconstituted in 0.5M, and following complete reconstitution, samples were diluted to 100mM NaOH, and total protein content was measured using the BCA kit.

The cell number/mg protein ratio was calculated on a separate occasion by manually counting cells through a hemocytometer (each sample was counted twice. For each count, four squares were analyzed). A known number of cells were lysed according to the Glutathione assay kit procedure described above. Deproteinized samples were reconstituted and analyzed as described above.

---

**Table 1.** List of antibodies used in this study

| Antibody | Supplier | Catalog | RRID | Dilution |
| --- | --- | --- | --- | --- |
| S100B | Abcam | ab52642 | AB_882426 | 1:100 |
| CD44 | Abcam | ab157107 | AB_2847859 | 1:500 |
| GFAP | DAKO | Z0334 | AB_10013382 | 1:500 |
| AQP4 | Sigma-Aldrich | HPA014784 | AB_1844967 | 1:125 |
| ALDH1L1 | Abcam | ab190298 | AB_2857848 | 1:500 |
| VIMENTIN | Abcam | ab92547 | AB_10562134 | 1:500 |
| EAAT1 | Thermo Fisher Scientific | PA5-19709 | AB_10982702 | 1:200 |
| EAAT2 | Thermo Fisher Scientific | 711020 | AB_2633106 | 1:250 |
| ICAM-1 | R&D Systems | BBA3 | AB_356950 | 1:50 |

**Table 2.** List of TaqMan probes used in this study.

| Gene | TaqMan Assay ID |
| --- | --- |
| ALDH1L1 | Hs01003842_m1 |
| CD44 | Hs01075864_m1 |
| DCX | Hs00167057_m1 |
| GAPDH | Hs02758991_g1 |
| GFAP | Hs00909233_m1 |
| NES | Hs04187831_g1 |
| NFIA | Hs00325656_m1 |
| S100B | Hs00902901_m1 |
| SLC1A2 | Hs01102423_m1 |
| SLC1A3 | Hs00904823_g1 |
| SOX1 | Hs01057642_s1 |
| SOX2 | Hs01053049_s1 |
| RELN | Hs01022646_m1 |
| SOX9 | Hs00165814_m1 |
| MAOB | Hs01106246_m1 |
| SLIT1 | Hs00171488_m1 |
| VIM | Hs05024057_m1 |
| GLUL | Hs01013055_g1 |
| AQP4 | Hs00242342_m1 |

